# Context-Aware Prediction of Pathogenicity of Missense Mutations Involved in Human Disease

**DOI:** 10.1101/103051

**Authors:** Christoph Feinauer, Martin Weigt

**Affiliations:** Sorbonne Universités, UPMC, Institut de Biologie Paris-Seine, CNRS, Laboratoire de Biologie Computationnelle et Quantitative UMR 7238, Paris, France

## Abstract

Amino-acid substitutions are implicated in a wide range of human diseases, many of which are lethal. Distinguishing such mutations from polymorphisms without significant effect on human health is a necessary step in understanding the etiology of such diseases. Computational methods can be used to select interesting mutations within a larger set, to corroborate experimental findings and to elucidate the cause of the deleterious effect. In this work, we show that taking into account the sequence context in which the mutation appears allows to improve the predictive and explanatory power of such methods. We present an unsupervised approach based on the direct-coupling analysis of homologous proteins. We show its capability to quantify mutations where methods without context dependence fail. We highlight cases where the context dependence is interpretable as functional or structural constraints and show that our simple and unsupervised method has an accuracy similar to state-of-the-art methods, including supervised ones.

## 1 Introduction

As of January 2017, the UniProt database (Magrane et al., 2011) lists 75.431 amino acid variants of their sequences and classifies 28.891 of them as associated with human diseases, many of which lethal. Since every human genome is estimated to contain 74 de novo SNVs and around 0.6 de novo missense mutations (Veltman and Brunner, 2012), the number of known variants and the need for their classification is likely to increase strongly in the near future due to cheap sequencing and the trend towards *personal genomics* (Angrist, 2016). The potential of the field can be appreciated considering private companies like 23andme, which routinely publish genome-wide association studies based on hundreds of thousands of human genomes (Chen et al., 2016; Okbay et al., 2016a, b; Strachan et al., 2016).

Problems arise when considering rare variants or experiments with small sample sizes, where the statistics remains insufficient to conclusively label a specific variant or a gene as disease-associated (Purcell et al., 2014). Another problem is that the sheer number of variants discovered prohibits a further experimental analysis of most of them. In both cases computational methods can help: They can be used for extracting interesting variants from a pool of unknown pathogenicity; and sometimes used to understand the etiology of the disease with which the variant is associated (Wang et al., 2012). Many computational methods for predicting pathogenicity of amino acid substitutions exist. Unsupervised methods look at different proxies for pathogenicity and are, for example, based on the frequencies of amino acids at the mutated residue in a multiple sequence alignment of homologous sequences (Kumar et al., 2009), or on the time of conservation of the mutated residue in a reconstructed phylogenetic tree (Tang and Thomas, 2016). Supervised methods combine such proxies and add other information like structural data or annotations (Adzhubei et al., 2013; Kircher et al., 2014). Unsurprisingly, they often perform better than unsupervised methods, but arguably not by a very large margin (Tang and Thomas, 2016; Grimm et al., 2015). Inspired by the successes of global probability models for protein sequences in fields like the prediction of protein residue contacts (Weigt et al., 2009; Jones et al., 2012; Morcos et al., 2011; Kamisetty et al., 2013), the inference of protein interaction networks (Ovchinnikov et al., 2014; Hopf et al., 2014; Feinauer et al., 2016) and the modeling of mutational landscapes in bacteria and viruses (Mann et al., 2014; Morcos et al., 2014; Figliuzzi et al., 2015), we propose in this work the inclusion of a new type of information when predicting the pathogenicity of mutations in humans: the sequence context in which the mutation appears. We first show that the sequence context indeed carries information useful in the prediction of pathogenicity by comparing the performance of two sequence-based models, one of which includes the context dependence and the other one not. We show cases in which this context dependence is interpretable as structural or functional constraints that the model exploits to correctly predict the pathogenicity of the mutation. In the age of machine learning, where results of algorithmss are often hard to interpret, this might be an important advantage of a method. We then go on and show that the performance of the method compares favorably with published methods, supervised and unsupervised.

## 2 Results

### Outline

We assess the pathogenicity of a mutation by first mapping the mutated residue to a consensus position in one of the profile hidden Markov models (pHMM) (Durbin et al., 1998) provided by the protein domain family database (Pfam) (Finn et al., 2016). We then fit a maximum entropy probability distribution to the multiple sequence alignment (MSA) of this protein domain family and calculate scores for pathogenicity based on this probability distribution. In order to assess whether the sequence context in which a mutation occurs carries information about its pathogenicity, we compare the predictive power of scores based on a context-dependent model with scores based on a context-independent model (short *independent model*). The scores reflect this context dependence or context-independence: Changing the sequence context in which the mutation occurs changes the score derived from the context-dependent model, while the score derived from the context-independent model remains the same.

### Independent Model

The independent model we use is a maximum-entropy distribution that reproduces amino acid frequencies in the MSA while ignoring covariances between amino acids. This model treats all residues in the MSA as independent and thus neglects any possible epistatic effects (see *Materials and Methods*). The probability distribution is the same as in profile models used for constructing MSAs, and for aligned positions in profile hidden Markov models when neglecting gaps (Durbin et al., 1998). Although such models cannot capture structural or functional constraints comprising more than one residue, they have a major advantage in terms of simplicity: The number of parameters scales linearly with the length of the sequences (see *Materials and Methods*), which means that relatively few sequences are needed to fit the model well.

### Context-Dependent Model

Context dependence can be included in our framework by using a probability distribution in which the residues in a protein sequence are not independent from each other. This means that the probability of finding an amino acid at some specific position depends on the amino acids found at other positions. Scores that quantify the pathogenicity of mutations based on such probability distributions show epistatic effects and the predicted pathogenicity of a mutation depends on the sequence context. In this work we use the direct-coupling analysis (DCA) (Weigt et al., 2009), which constructs a maximum-entropy model constrained to reproduce the covariances between amino acids in the MSA. It includes context dependence and has been used successfully for modeling protein sequences (see *Materials and Methods* and Stein et al., 2015 for a general introduction). The inference of the model follows Ekeberg et al., 2013. The context-dependent model should be able to capture structural or functional constraints comprising more than one residue, but its parameters scale quadratically with the length of the protein (see *Materials and Methods*). We show below that the available sequences are nonetheless sufficient to make the context-dependent model perform better than the independent model. A feature of the context-dependent model is that the influence of the sequence context is easily analyzed: Part of the change in log probability when substituting an amino acid in a sequence is made up of contributions from other residues. We call these contributions *c*_*j*_, where *j* labels a non-mutated residue. We show below that the *c*_*j*_ can be used to interpret the score (see *Materials and Methods*).

### Scoring Pathogenicity

A natural score for the pathogenicity of a mutation is the difference in the logarithm of the probabilities of the mutated and the original sequence (Figliuzzi et al., 2015). We call this score Δ*L*. We show below that this quantity gives good results, but is not optimal. This is probably due a dependence of the expected value of the difference in the logarithms on the length of the protein domain and the sampling depth of the original MSA. This makes a comparison of this score across different protein domain families problematic. We therefore propose another measure *r* that is based on a imaginary mutagenesis experiment: We calculate Δ*L* for all possible single-site amino acid substitutions in the original sequence and determine which rank the mutated sequence to be assessed has in this spectrum. We then map this rank to the interval between 0 and 1 by dividing by the number of different Δ*L* in the hypothetical experiment. We show below that this score outperforms Δ*L* significantly. We did not attempt to define thresholds for classification of mutations as deleterious or benign. Such a classification should be addressed in a supervised framework using r as an additional input. This we reserve for future investigations. Throughout the rest of the paper, we call *r* and Δ*L* the scores calculated on the context-dependent model, and *r*^*IM*^ and Δ*L*^*IM*^ the scores calculated on the independent model.

### Comparison between Context-Dependent Model and Independent Model

We calculated the scores *r*,*r*^*IM*^, Δ*L* and Δ^*IM*^ for the *SwissVarSelected* (Grimm et al., 2015) benchmark set. Of the 11028 mutations in this dataset that we could map to a UniProt accession number, about 49% could be mapped to a consensus position in a Pfam domain family. We calculated the receiver operating characteristic curve (Friedman et al., 2001) for the various scores and show them them in Fig. 1. We first notice that the scores based on imaginary mutagenesis experiments, *r* and *r*^*IM*^ perform better than their counterparts Δ*L* and Δ*L*^*IM*^. We furthermore notice that the scores derived from context-dependent models, *r* and Δ*L* outperform the scores derived from independent models, *r*^*IM*^ and Δ*L*^*IM*^. The best-performing score is *r*. We therefore conclude that context dependence represents indeed additional and useful information for the prediction of pathogenicity. To further corroborate these results we repeated the test on all annotated mutations and polymorphisms in the Uniprot Database (Magrane et al., 2011). While the overall performance of all models on this dataset is higher than on the *SwissVarSelected* dataset (in line with Tang and Thomas, 2016), the relative performances remain approximately equal (see Fig. 1).

### Sampling Depth

Since the context-dependent model is based on a large number of parameters (see *Materials and Methods*), it is important to have a sufficient number of homologous sequences in the model inference process. We used the *effective number of sequences* in the MSA as calculated in the inference process (see Ekeberg et al., 2013) to estimate the amount of information in the MSA. This differs from the number of sequences in the alignment because of corrections for phylogenetic and experimental biases (Ekeberg et al., 2013). The mean number of effective sequences for the mutations in the *SwissVarSelected* that could be mapped to Pfam families is 9018 sequences, with a median of 1923 (a histogram of the number of effective sequences can be found in Fig. 2). While this average would be considered as sufficient for structural predictions (Morcos et al., 2011), we also observe a large number of mutations that correspond to multiple sequence alignments with less than 500 effective sequences. This would be considered as not sufficient in structural predictions. Since is not clear how the performance in structural prediction relates to the performance in the prediction of pathogenicity, we quantified the effect of sampling depth on predictive performance. We divided the *SwissVarSelected* dataset in three approximately equal parts by the number of effective sequences in the corresponding Pfam families. The corresponding thresholds were 754 and 5828 effective sequences. We then calculated the AUROCs for the context-dependent model and the independent model after sampling from each subset to reach the same proportion of pathogenic and non-pathogenic variants as in the original dataset (approximately 40% to 60%). To our surprise, the difference in performance when using *r* as a score was not very pronounced in the three subsets (see Fig. 5 in the *Supplemental Material*). We take this as evidence that even for the tercile with the least sequences the number of sequences is sufficient. We also observed that for the largest tercile the performance worsens and for the independent model the performance seems to generally deteriorate with higher sequences numbers (see 5 in the *Supplemental Material*). This could be either due to systematic problems with the largest families, like false labeling or non-conservative inclusion thresholds of the MSAs, or to random variability in the data set. A further partitioning of the smallest tercile shows that the performance with between 139 and 346 sequences is only marginally worse than with more than 346 sequences (see Fig. 7 the *Supplemental Material*). A minimum of 139 sequences includes 89% of the mutations in the test set (see Fig. 2). The general outlook seems therefore to be that for most mutations that can be mapped to a position in a PFAM domain the number of sequences is sufficient. The problem that many mutations cannot be mapped to a Pfam domain (about 51% in the *SwissVarSelected* dataset) is likely to pose a larger challenge in actual usage than lacking sampling depth.

**Figure 1:**
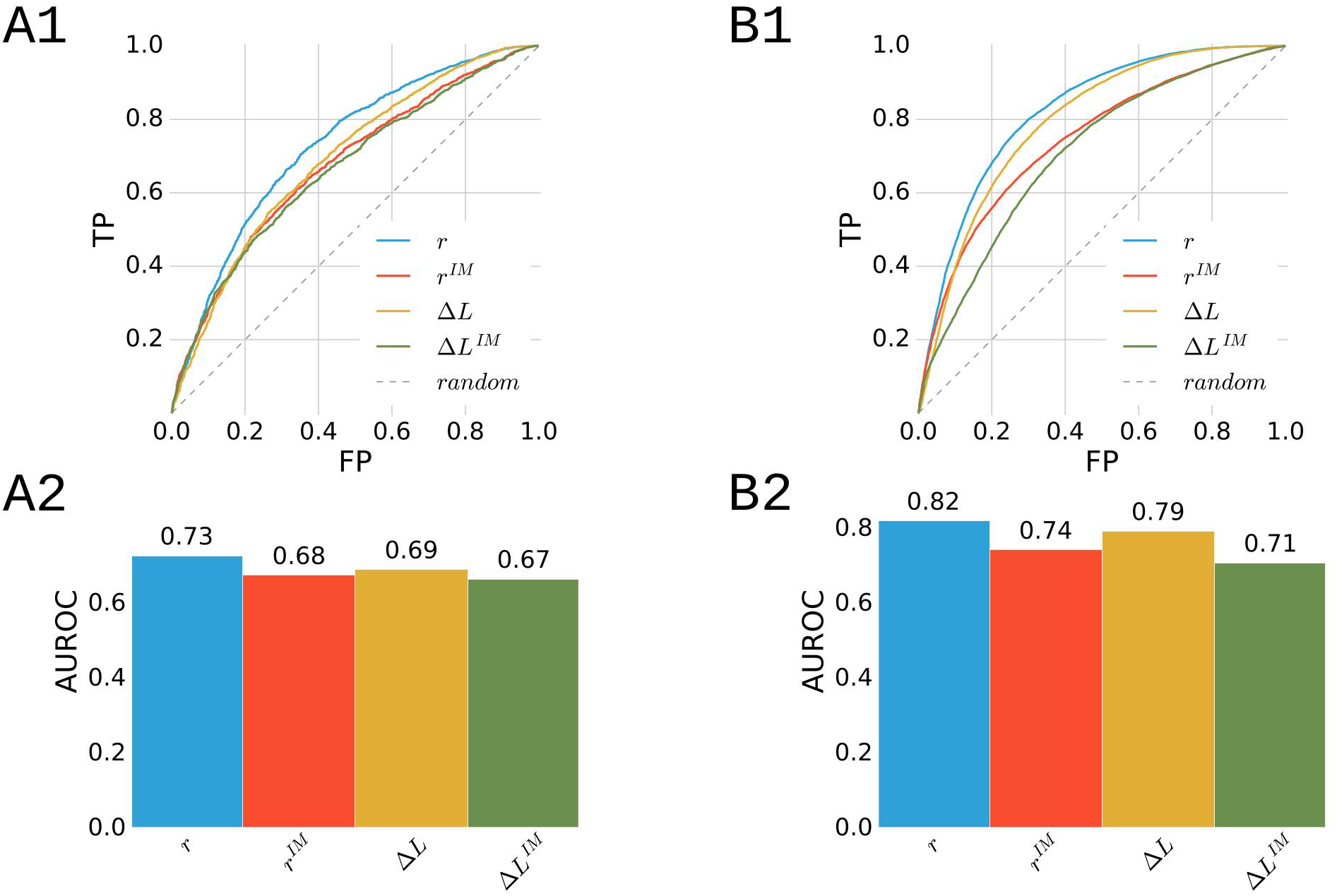
**A1,A2:** ROC curves (A1) and area under the curves (A2) for scores from the context-dependent (Δ*L* and *r*) and the independent model (Δ*L*^*IM*^ and *r*^*IM*^), calculated on the *SwissVarSelected* (Grimm et al., 2015) dataset where the method returned predictions. TP is true positive rate and FP false positive rate. **B1,B2:** ROC curves (B1) and area under the curves (B2) for scores from the context-dependent (Δ*L* and *r*) and the independent model (Δ*L*^*IM*^ and *r*^*IM*^), calculated on all mutations annotated in UniProt (Magrane et al., 2011) where the method returned predictions. TP is true positive rate and FP false positive rate.

### Comparison with other Methods

The central message of this work is that context dependence is important for predicting the pathogenicity of amino acid substitutions. It is nonetheless interesting to compare the performance of our method to state-of-the-art predictors. As one would expect the best performing methods today are supervised methods that take information from different sources like sequence data or experimental structures into account (Grimm et al., 2015). In Fig. 3 we compare our context-dependent model and the independent model to: Panther-PSEP, a recent unsupervised method based on phylogeny (Tang and Thomas, 2016); SIFT, a highly cited unsupervised method using residue conservation as a measure for pathogenicity (Kumar et al., 2009); Polyphen-2, a supervised method using 11 features derived from homologous sequences and structural data (Adzhubei et al., 2013); CADD, a supervised method for genetic variants using 63 annotations as inputs, including protein level annotations derived from other pathogenicity-predictors like Polyphen or SIFT (Kircher et al., 2014). In Fig. 3 we show the AUROC values achieved by the different methods. All methods have AUROCs around 0.7, with supervised methods slightly outperforming the unsupervised methods. It is surprising that the large number of features (including structural features) that the supervised methods exploit do not lead to a larger boost in performance, but this has been observed before (Thusberg et al., 2011). While the unsupervised methods generally show a very similar performance, we observe by comparing the performance of *r* and *r*^*IM*^ that the inclusion of the context dependence lifts the performance of our model from the worst-performing unsupervised method to the best-performing unsupervised method presented. As a last point we notice that the performance of PANTHER in our runs is at odds with the performance reported in Tang and Thomas, 2016 (area under the curve 0.72 vs. 0.70). We speculate this to be a data artifact: Since we had several methods in the comparison, we reduced the *SwissVarSelected* dataset to only the mutations for which all methods returned results. This left only 3857 of 12739 mutations; and since the authors in Tang and Thomas, 2016 presumably did the same, the overlap between the two datasets used might be even smaller.

**Figure 2:**
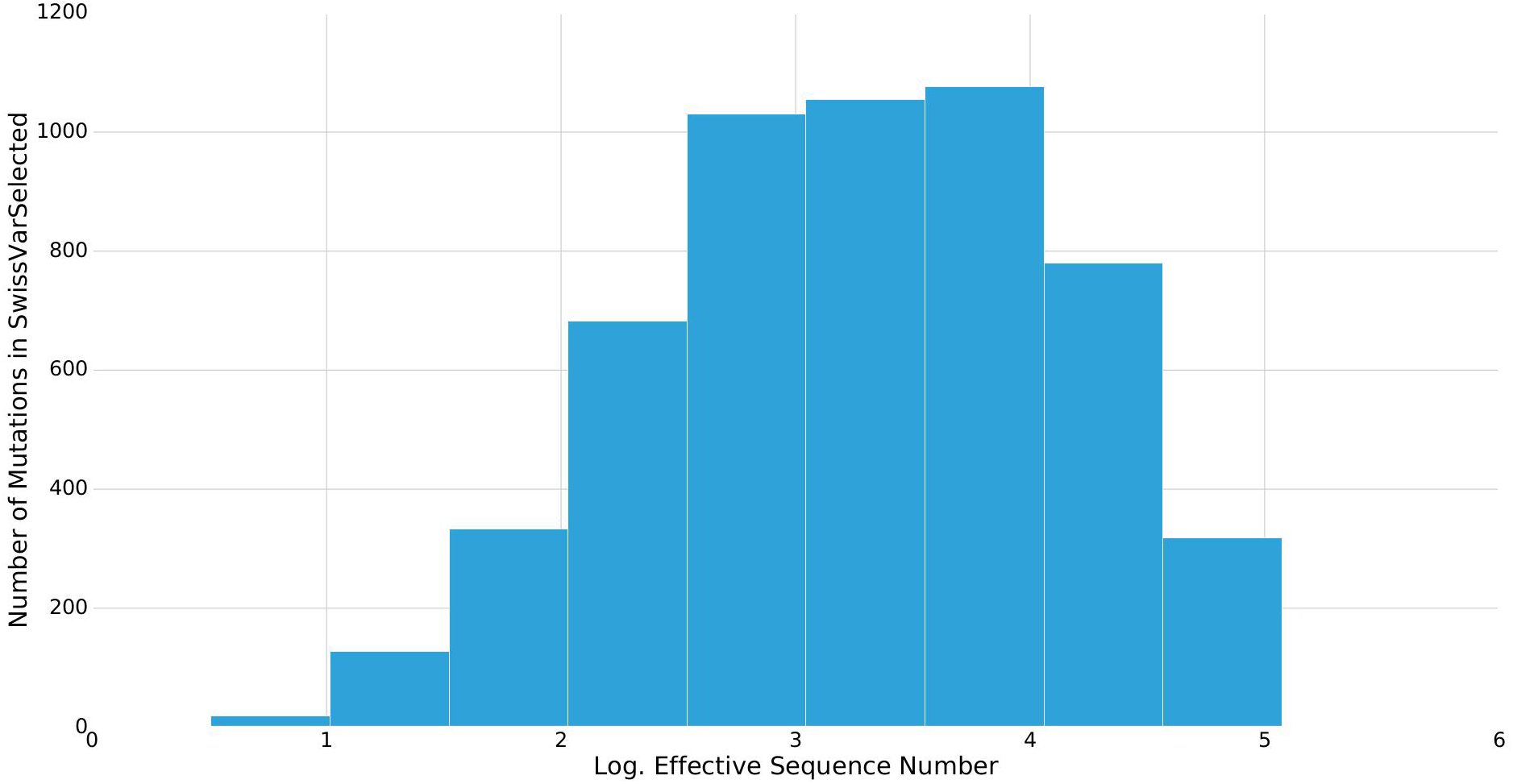
Histogram of the decadic logarithm of the effective sequence numbers in the multiple sequence alignments corresponding to the mutations in the *SwissVarSelected* dataset (Grimm et al., 2015). Only mutations that could be mapped to a Pfam family are included.

## 3 Discussion

In this work, we have shown that including context dependence in methods for predicting the pathogenicity of amino acid substitutions can lead to a significant boost in performance. We have also shown that a relatively simple unsupervised method based on maximum entropy modeling and using no biological prior information can compete with state-of-the-art unsupervised and supervised methods.

One of the characteristics of the probabilistic model constructed by DCA is the explicit estimation of epistatic couplings between different residues. This allows the disentangling of the context dependence into contributions from individual residues in the unmutated sequence, and to **interpret context dependence** in a number of sample cases. In this respect, we speculate that the context dependence helps predicting the pathogenicity of mutations because the corresponding probability distribution captures constraints acting on groups of residues, i.e. residues that are part of a functional group or form structural bonds with each other. In order to validate this point, we took a closer look at mutations that were labeled as disease-associated in the dataset, had a high *r*-score in the context-dependent model and a low *r*^*IM*^ score in the independent model. We found 40 pathogenic mutations for which *r*-*r*^*IM*^ > 0.5. We then checked existing literature on these mutations for examples in which our models could be interpreted consistently with this literature. We found evidence for at least three cases in which context dependence can lead to better performance. The three cases are 1) mutations within functional groups, 2) mutations leading to a loss of a structural contact, 3) mutations that are common in one cluster of sequences appearing in another cluster. We elaborate these cases with examples.

**Figure 3:**
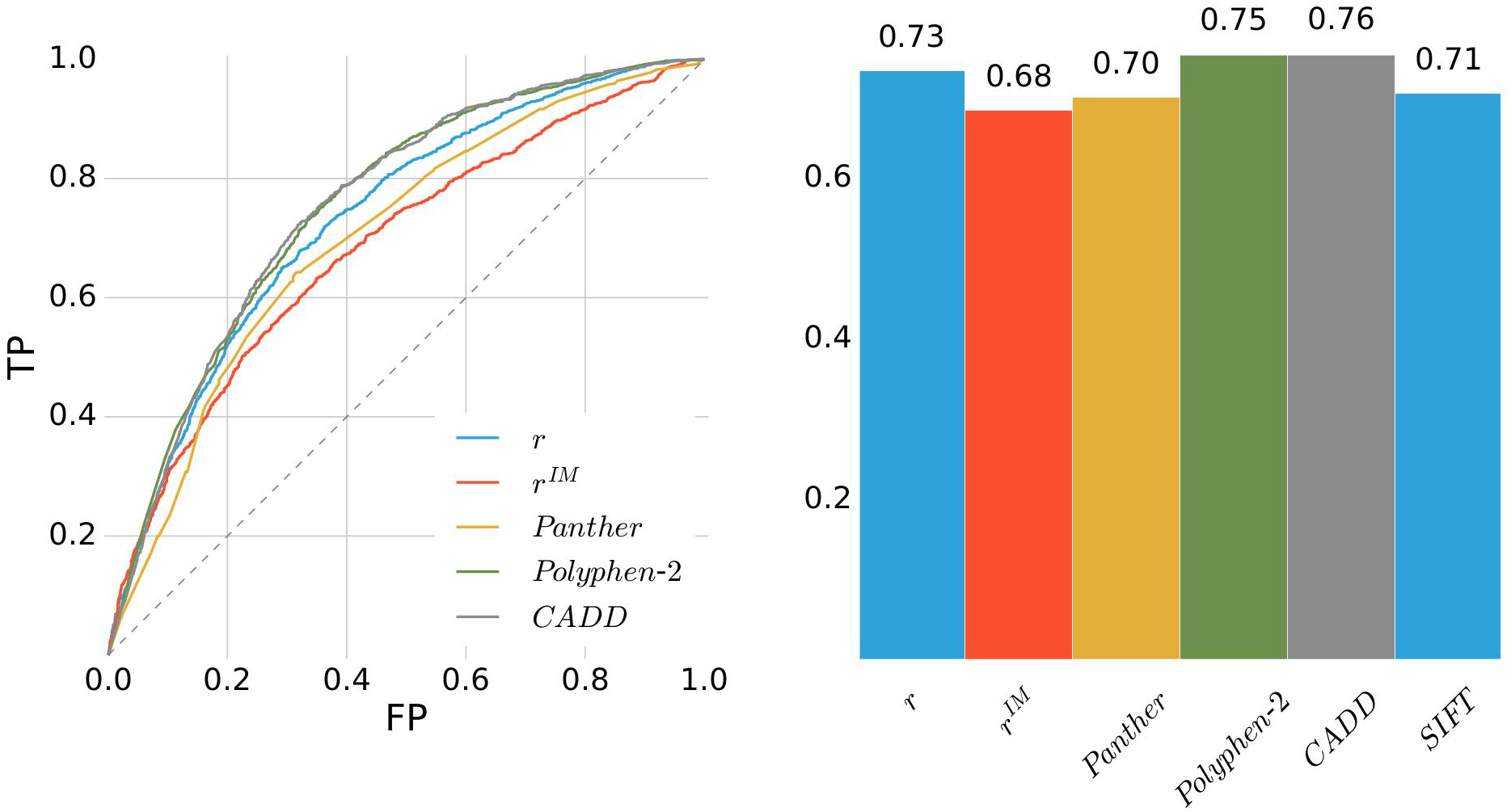
ROC curves (left) and area under the curves (right) for scores from the context-dependent model (*r*) and the independent model (*r*^*IM*^) in comparison with other methods as labeled. The results for the other methods were obtained as described in *Materials in Methods*. The dataset used here is *SwissVarSelected* (Grimm et al., 2015) and only mutations where kept where all methods returned predictions. TP is true positive rate and FP is false positive rate.

### Adrenal Cushing’s Syndrome

Patients with adrenal Cushing’s syndrome exhibit a variety of symptoms stemming from an overexposure to cortisol (Goh et al., 2014). A specific mutation that has been found in such tumors is PRKACA^L206R^, affecting a subunit of a protein kinase. This mutation prevents that PRKACA is bound by its regulator PRKAR1A, increasing its phosphorylation activity and leading to cortisol production downstream (Goh et al., 2014). Since both arginine and leucine are common at this position in the corresponding protein domain family, the independent model assigns both sequences similar rank (*r*^*IM*^ = 0.14 for PRKACA and *r*^*IM*^ = 0.153 for PRKACA^L206R^. In contrast, the context-dependent model scores the mutated sequence considerably worse than the original sequence (*r* = 0.02 for PRKACA and *r* = 0.73 for PRKACA^L206R^). In order to understand this, we calculated the contributions from other residues to the corresponding drop in Δ*L* in the context-dependent model when exchanging the arginine with the leucine (see *Materials and Methods*). This contribution quantifies how strongly the specific amino acid found in some other position favors or disfavors the mutation. In Fig. 4A we color residue 206 in blue, the 4 strongest contributors in yellow and the regulator of PRKACA in red. Of these 4 only two, 201G and 200C, qualify as contacts with 3.7 Å and 3.2 Å minimal heavy atom distance to 206L (distances are extracted from the PDB 3TNP (Zhang et al., 2012) of a mouse homolog of the human protein). The other two, 197W and 238P, are more distant on the chain and also more distant in the structure with 10 Å and 8.9 Å minimal heavy atom distance to 206L. However, all four are in the interaction surface between PRKACA and an 11-residue segment (95R to 106T) of its regulator PRKAR1A, with 21 contacts between this segment and the 4 contributors and 206L. We take this as evidence that the context-dependent model has captured a multi-residue constraint acting on residues within the functional group responsible for the interaction. This functional group represents the sequence context that allows the correct classification of this mutations as pathogenic. This is in full agreement with the idea that the mutation impairs the binding of the regulator PRKAR1A to PRKACA (Goh et al., 2014).

### Classic Maple Syrup Urine Disease

The disease is caused by an impaired metabolism of branched-chain amino acids (Chace et al., 1995). Carriers often die young, within months or days after birth (Wang et al., 2012). The mutation BCKDHB^Q346R^ has been implicated in the disease (Wang et al., 2012). The BCKDHB gene encodes the two β-subunits of the *E*1-complex, which is part of the multi-subunit enzyme complex BCKD involved in the metabolism of leucine, isoleucine and valine. (Indo et al., 1987). There is evidence that the mutation destabilizes the interaction of 346R with 357I between the βsubunits (Wang et al., 2012). The two residues are 2.72 Å apart in the complex (minimum distance between heavy atoms, PDB 1X7Y (Wynn et al., 2004)). The rank of the mutated sequence is considerably worse in the context-dependent model (*r* = 0.83) than in the independent model (*r*^*IM*^ = 0.27). We speculated that this might be due to a structural constraint between residue 346 and residue 357 that the independent model cannot capture. Shifting to structural analysis, we calculated *F*^*APC*^ scores for the residues (see *Materials and Methods*). These are quantities determined by the model that can be used to infer residue contacts (Ekeberg et al., 2013). We found that the residue with the strongest coupling to residue 346 was indeed residue 357 (excluding short-range couplings), indicating a strong interaction between the residues in the context-dependent model (see Fig.4B). We therefore conclude that the context-dependent model captured a structural constraint involved in the homo-dimerization of the two β-subunits. The sequence context that helped the context-dependent model to asses pathogenicity in this interpretation is reduced to a single residue.

**Figure 4:**
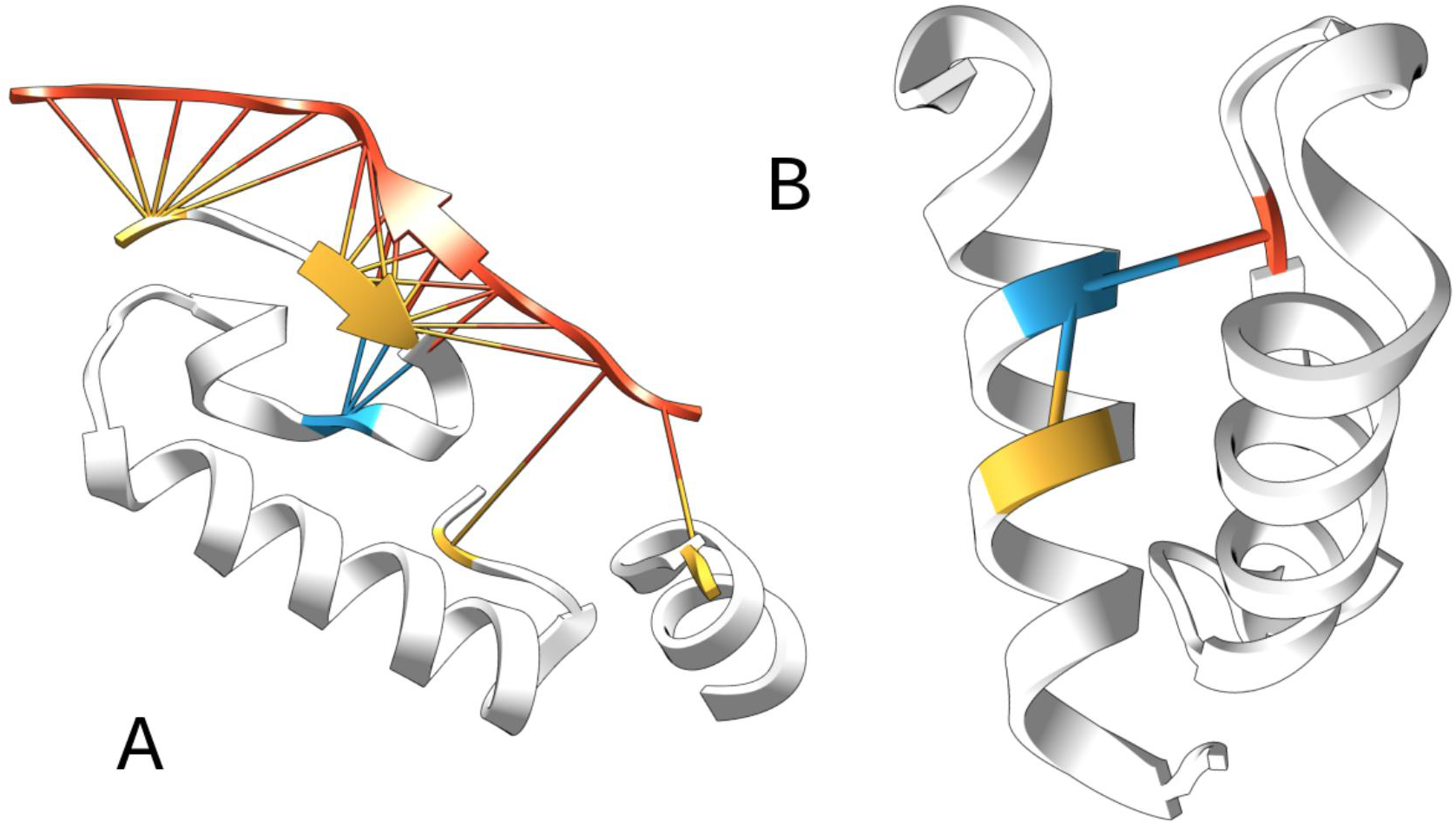
**A** Interaction between PRKACA (lower left) and its regulator PRKAR1A (upper right, red). The yellow residues are the strongest contributors to the drop in Δ*L* when exchanging an arginine with a leucine at residue 206 of PRKACA (blue). The sticks represent residue contacts between PRKACA and its regulator. The PDB used is 3TNP (Zhang et al., 2012). This figure shows only fragments for clarity. See *Supplemental Material* for the full structure. **B** Interaction between two *E*1-β-subunits within the BKCD complex. Sticks represent the two strongest contacts by *F*^*APC*^ score of residue 346 (blue). Only residue 346 of the left subunit and its contacts are colored: an α-helix intra-chain contact with residue 342 (yellow) and a homomeric contact with residue 357 of the right chain (red). A trivial contact with residue 347 is not shown. The PDB used is 1X7Y (Wynn et al., 2004). This figure shows only fragments for clarity. See *Supplemental Material* for the full structure.

### Alpha-Mannosidosis

The disesae is caused by a deficient catabolism of N-linked oligosaccharides, due to a defective enzyme lysosomal alpha-mannosidase encoded in the gene MAN2B1. Patients might suffer from symptoms like mental retardation, changed facial features and impaired hearing abilities (Roces et al., 2004). A mutation implicated in the disease is MAN2B1^R202P^ (Stensland et al., 2012). It has a high pathogenicity-score in the context-dependent model (*r* = 0.89), but a low score in the independent model (*r*^*IM*^ = 0.14). Calculating the largest contributors (see *Materials and Methods*) to the change in the score, we find contributions of similar magnitude from many residues. An analysis of PDB 1O7D (Heikinheimo et al., 2003) revealed that only 4 of the ten largest contributors are in contact with residue 202. Furthermore, these 4 are all less than 8 residues apart from it. A possible explanation for such cases is that the sequences cluster and the mutation exchanges an amino acid common at the residue in one cluster with an amino acid common in another cluster. The context-dependent model might not capture biologically significant constraints in such cases, but can nonetheless single out residues that are uncommon in the specific context. In fact, we observe that sequences that have an arginine at residue 202 are significantly more similar to each other (mean Hamming distance 0.51) than to sequences that have a proline at that residue (mean Hamming distance 0.74). This can be summed up by saying that the sequence context that helped the context-dependent model to assess the pathogenicity in this case is the entire protein sequence. This is of course a hypothesis of last resort. Another plausible explanation is that there is biological constraint captured by the context-dependent model, but we are not able to discover it.

### Note

While finalizing the redaction of this article, another article with partially overlapping results (Hopf et al., 2017) has been published online. The paper comes to a similar conclusion, i.e. the inclusion of sequence context dependencies improves the prediction of mutation effects. While a direct comparison is difficult since the paper uses different data sets, it is built upon the Δ*L* log-probability score, which on all our data-sets performs significantly worse than the *r* rank score proposed in our work. We also note that our main data-set, *SwissVarSelected*, was explicitly designed as a comparable benchmark and used as such in recent publications (Grimm et al., 2015; Tang and Thomas, 2016). We therefore believe that our work gives additional and valuable insights on how modeling context dependence affects predictive performance compared to established methods.

## 4 Materials and Methods

### Outline of the Method

A mutation is assumed to be given as a Uniprot Accession Number, a position, a reference amino acid and a substituted amino acid. The Pfam domain architecture of the corresponding Uniprot sequence is extracted from the Pfam database (Finn et al., 2016) and it is checked whether the mutated residue can be mapped to a consensus column of a Pfam pHMM. If not, the method does not return a result. If yes, plmDCA is run on the Pfam MSA of the protein domain family and the various scores are calculated with its output. For the independent model the inference with plmDCA is replaced by a analytic formula.

### Context-Dependent Model

The context-dependent model defines a maximum entropy probability distribution *p*(*a*) for all amino acid sequences of length *N* (Morcos et al., 2011). The model can be derived by enforcing the constraint that it should reproduce the covariances between amino acids in the MSA (Stein et al., 2015). The logarithm of this distribution can be written as

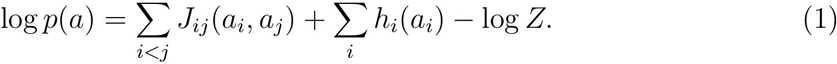

Here, *a*_*i*_ is the amino acid at residue *i*, the fields *h*_*i*_(*a*) represent the propensity of position *i* to be amino acid *a* and *J*_*ij*_(*a, b*) are coupling parameters that represent the context dependence in the model. log *Z* is a normalization constant. The parameters are inferred using plmDCA as described in (Ekeberg et al., 2013).

An intuitive score for the pathogenicity of substituting amino acid *a*_*i*_ with *a*^*′*^_*i*_ in sequence *a* is the change in log probability (Figliuzzi et al., 2015)

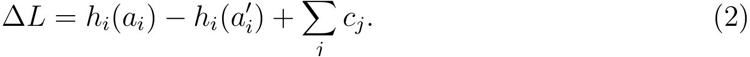

The *c*_*j*_ measure the contribution of residue *j* to the score,

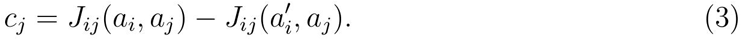

However, the model for a protein domain of even medium length *N* = 50 has already more than 5×10^5^ coupling parameters, which are often inferred on MSAs of 1000 sequences or less. We expect this measure therefore to be very noisy.

A general measure for the interaction between *i* and *j* are the *F*^*APC*^ scores used in structure prediction, which can be calculated from the *J*_*ij*_ (Ekeberg et al., 2013). They contain a sum over many couplings and we expect them to be more stable than the *c*_*j*_. Since the *F*^*APC*^ scores have been shown to capture structural contacts, we expect them to be useful when the substitution of an amino acid violates as structural constraint.

Another measure for the pathogenicity of a mutation is the rank of the mutated amino acid sequence in a hypothetical mutagenesis experiment. To this end, we calculated Δ*L* for all possible single amino acid substitutions in the original sequence. We then divided the rank of the mutated sequence in this list with the number of unique Δ*L* values in the hypothetical experiment. The resulting score r lies between 0 and 1 and is independent of the scale of Δ*L*.

### Independent Model

The independent model defines another maximum entropy probability distribution. It can be derived by enforcing the constraint that the model should reproduce the amino acid frequencies in the MSA. It does not contain any context dependence. The log probability can be written as

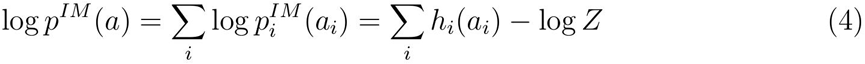

Using the same regularization (with regularization parameter λ set to 0.01) and reweighting techniques as in Ekeberg et al., 2013, an analytical solution can be found to be

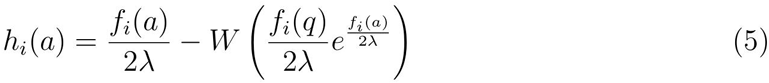

where *f*_*i*_(*a*) is the reweighted frequency of amino acid *a* at position *i*, *q* the most frequent amino acid at *i* and *W* the Lambert W function. We define Δ*L*^*IM*^ and *r*^*IM*^ analogously to above.

### Datasets and other Methods

The dataset *SwissVarSelected* has recently been shown to be a good benchmark set for the prediction of pathogenicity (Grimm et al., 2015). We used it as the main dataset since it does not overlap with the training sets of the supervised methods with which we compare our method. The dataset and the results of the methods SIFT (Ng and Henikoff, 2003) and CADD (Kircher et al., 2014) have been taken from (Grimm et al., 2015). Polyphen-2 (Adzhubei et al., 2013) has been run on the webserver provided by the authors. Panther (Tang and Thomas, 2016) has been run with the standalone software provided by the authors. As an additional dataset we downloaded all annotated Uniprot mutations from here. Pfam version 30.0 was used (Finn et al., 2016).

## Acknowledgments

We are grateful to Matteo Figliuzzi and Olivier Tenaillon for many discussions. MW acknowledges funding by the ANR project COEVSTAT (ANR-13-BS04-0012-01).

## Additional information

The author(s) declare no competing financial interests.

## 5 Supplemental Material

**Figure 5:**
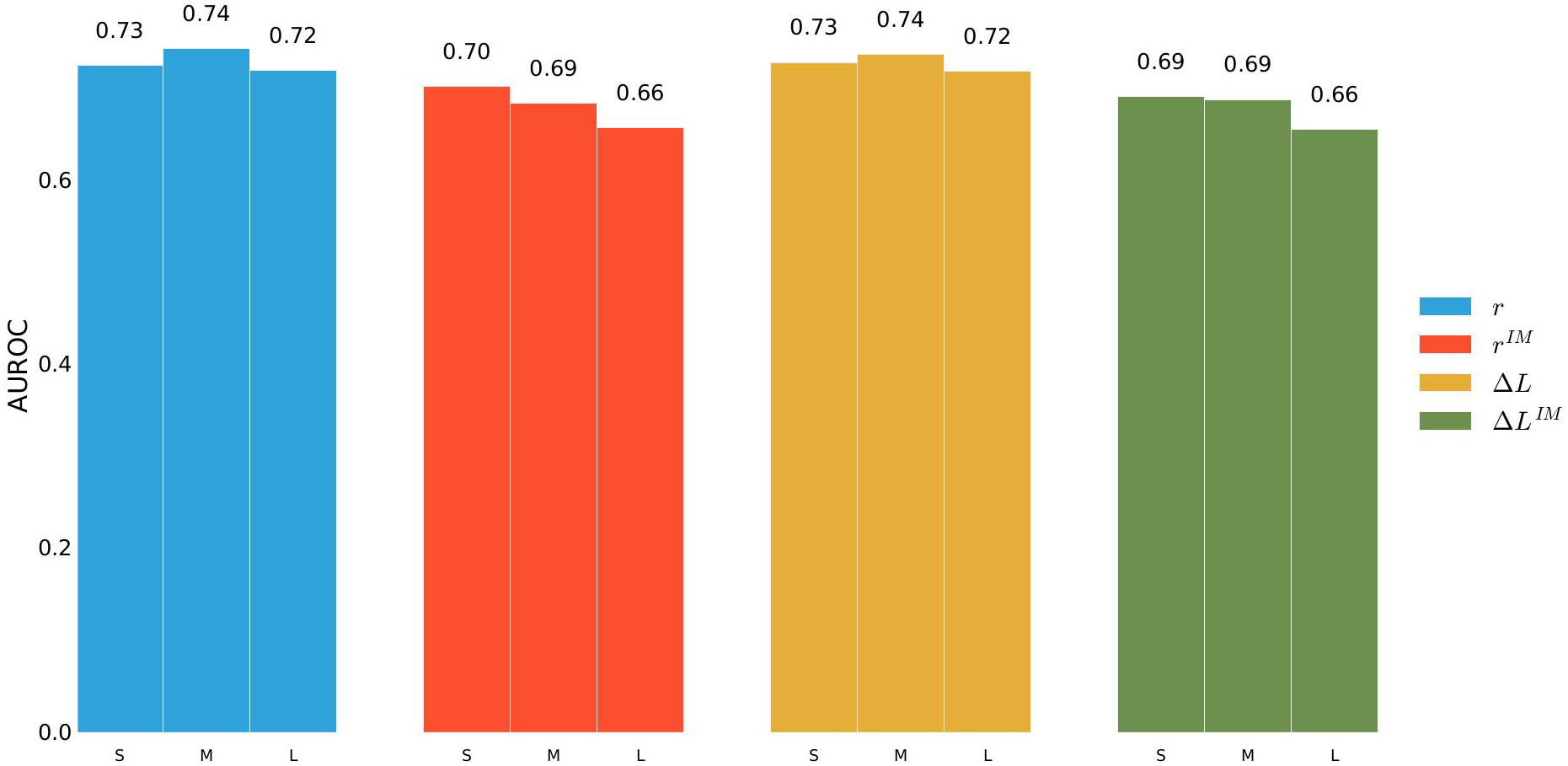
Performance (AUROC) of *r* score on subsets of *SwissVarSelected*. Subsets are labeled S (mutations with effective number of sequences less than 754), M (between 754 and 5828 effective sequences) and L (more than 5228 effective sequences). To evaluate performance the subsets were downsampled to arrive at the same fraction of pathogenic/non-pathogenic mutations (40% to 60%) as in *SwissVarSelected*.

**Figure 6:**
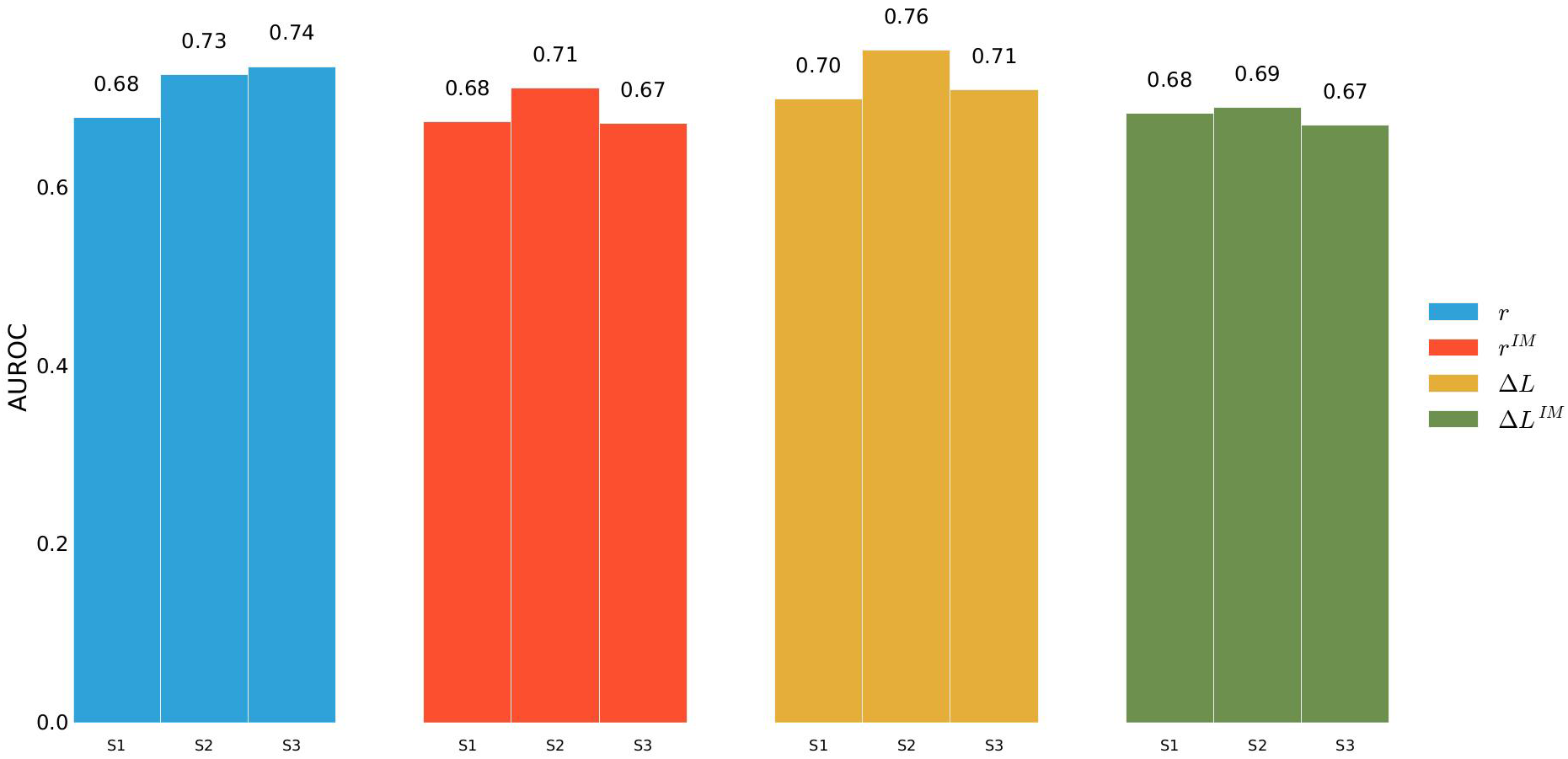
Bounds are: S1 0-139, S2 139-346, S3 346-749

**Figure 7:** Performance (AUROC) of *r* score on subsets of *SwissVarSelected*. Subsets are labeled S (mutations with effective number of sequences less than 139), M (between 139 and 346 effective sequences) and L (between 346 and 749 sequences). To evaluate performance the subsets were downsampled to arrive at the same fraction of pathogenic/non-pathogenic mutations (40% to 60%) as in *SwissVarSelected*.

**Figure 8:**
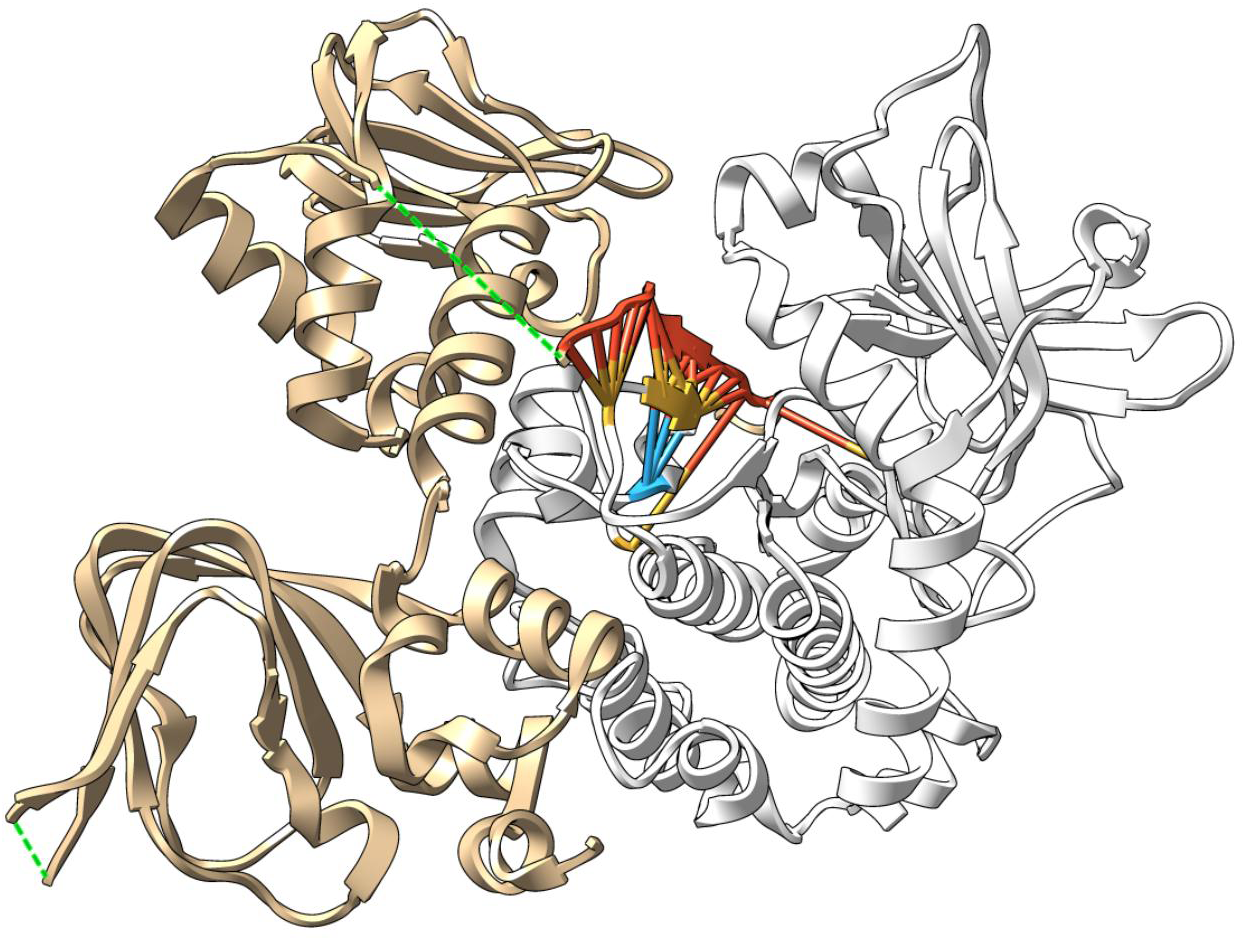
Interaction between PRKACA (lower left) and its regulator PRKAR1A (upper right, red). The yellow residues are the strongest contributors to the drop in Δ*L* when exchanging an arginine with a leucine at residue 206 of PRKACA (blue). The sticks represent residue contacts between PRKACA and its regulator. The PDB used is 3TNP (Zhang et al., 2012).

**Figure 9:**
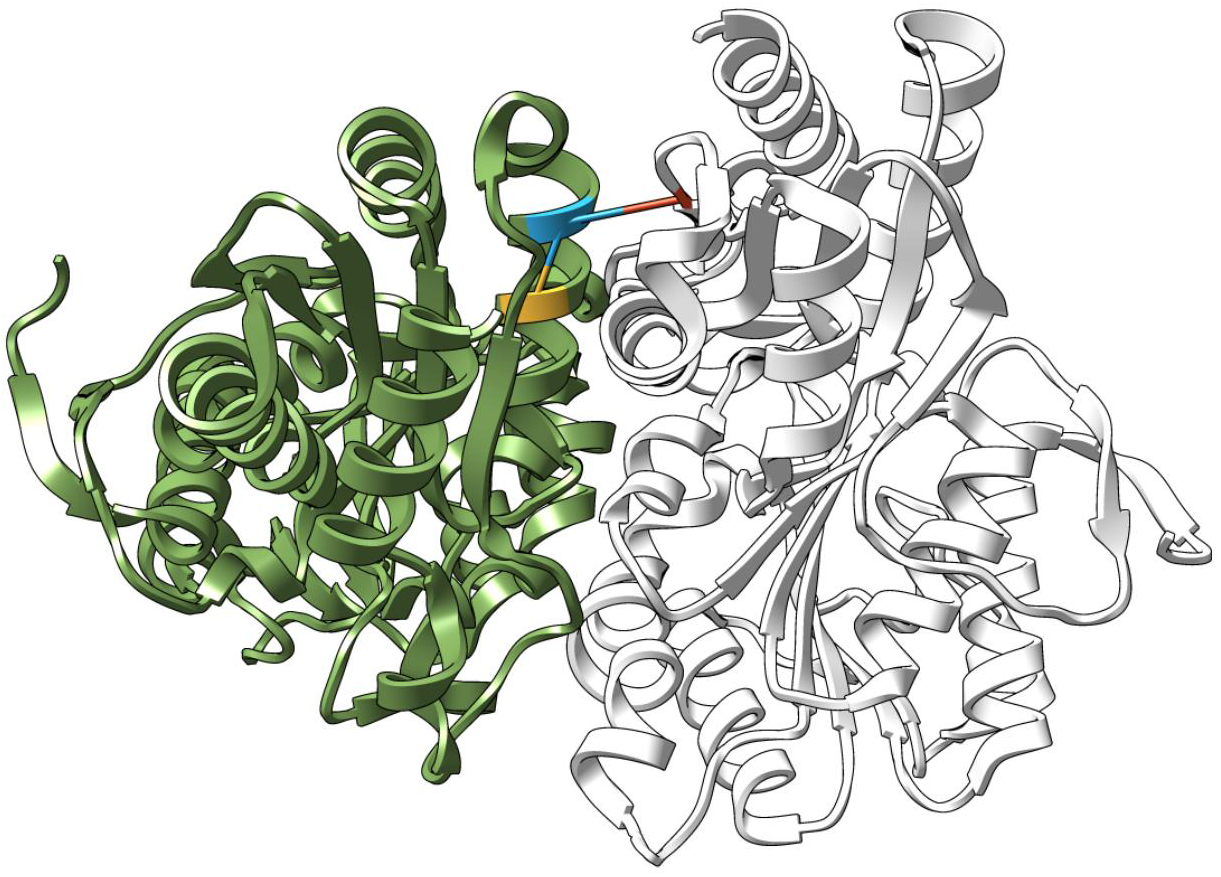
Interaction between two E1-β-subunits (green and white) within the BKCD complex. Sticks represent the two strongest contacts by *F*^*APC*^ score of residue 346 (blue). Only residue 346 of the left subunit and its contacts are colored: an α-helix intra-chain contact with residue 342 (yellow) and a homomeric contact with residue 357 of the right chain (red). A trivial contact with residue 347 is not shown. The PDB used is 1X7Y (Wynn et al., 2004).

